# Machine learning algorithms for systematic review: reducing workload in a preclinical review of animal studies and reducing human screening error

**DOI:** 10.1101/255760

**Authors:** Alexandra Bannach-Brown, Piotr Przybyła, James Thomas, Andrew S.C. Rice, Sophia Ananiadou, Jing Liao, Malcolm Robert Macleod

## Abstract

**Background:** Here we outline a method of applying existing machine learning (ML) approaches to aid citation screening in an on-going broad and shallow systematic review of preclinical animal studies, with the aim of achieving a high performing algorithm comparable to human screening.

**Methods:** We applied ML approaches to a broad systematic review of animal models of depression at the citation screening stage. We tested two independently developed ML approaches which used different classification models and feature sets. We recorded the performance of the ML approaches on an unseen validation set of papers using sensitivity, specificity and accuracy. We aimed to achieve 95% sensitivity and to maximise specificity. The classification model providing the most accurate predictions was applied to the remaining unseen records in the dataset and will be used in the next stage of the preclinical biomedical sciences systematic review. We used a cross validation technique to assign ML inclusion likelihood scores to the human screened records, to identify potential errors made during the human screening process (error analysis).

**Results:** ML approaches reached 98.7% sensitivity based on learning from a training set of 5749 records, with an inclusion prevalence of 13.2%. The highest level of specificity reached was 86%. Performance was assessed on an independent validation dataset. Human errors in the training and validation sets were successfully identified using assigned the inclusion likelihood from the ML model to highlight discrepancies. Training the ML algorithm on the corrected dataset improved the specificity of the algorithm without compromising sensitivity. Error analysis correction leads to a 3% improvement in sensitivity and specificity, which increases precision and accuracy of the ML algorithm.

**Conclusions:** This work has confirmed the performance and application of ML algorithms for screening in systematic reviews of preclinical animal studies. It has highlighted the novel use of ML algorithms to identify human error. This needs to be confirmed in other reviews, , but represents a promising approach to integrating human decisions and automation in systematic review methodology.

## Background

The rate of publication of primary research is increasing exponentially within biomedicine [1]. Researchers find it increasingly difficult to keep up with new findings and discoveries even within a single biomedical domain, an issue that has been emerging for a number of years [2]. Synthesising research – either informally or through systematic reviews - becomes increasingly resource intensive as searches retrieve larger and larger corpuses of potentially relevant papers for reviewers to screen for relevance to the research question at hand.

This increase in rate of publication is seen in the animal literature. In an update to a systematic review of animal models of neuropathic pain, 11,880 further unique records were retrieved in 2015, to add to 33,184 unique records identified in a search conducted in 2012. In the field of animal models of depression, the number of unique records retrieved from a systematic search increased from 70,365 in May 2016 to 76,679 in August 2017.

The use of text-mining tools and machine learning (ML) algorithms to aid systematic review is becoming an increasingly popular approach to reduce human burden and monetary resources required and to reduce the time taken to complete such reviews [3; 4; 5]. ML algorithms are primarily employed at the screening stage in the systematic review process. This screening stage involves categorising records identified from the search into ‘Relevant’ or ‘Not-Relevant’ to the research question, typically performed by two independent human reviewers with discrepancies reconciled by a third. This decision is typically made on the basis of the title and abstract of an article in the first instance. In previous experience at CAMARADES (Collaborative Approach to Meta-Analysis and Review of Animal Data from Experimental Studies), screening a preclinical systematic review with 33,184 unique search results took 9 months, representing (because of dual screening) around 18 person months in total. Based partly on this, we estimate that a systematic review with roughly 10,000 publications retrieved takes a minimum of 40 weeks. In clinical systematic reviews, Borah and colleagues [6] showed the average clinical systematic review registered on PROSPERO (International Prospective Register of Systematic Reviews) takes an average 67.3 weeks to complete. ML algorithms can be employed to learn this categorisation ability, based on training instances that have been screened by human reviewers [7].

Several applications of ML are possible. The least burdensome is when a review is being updated, where categorisations from the original review are used to train a classifier, which is then applied to new documents identified in the updated search [7; 8; 9]. When a screening is performed de novo, without such previous collection, humans first categorise an initial set of search returns, which are used to train an ML model. The performance of the model is then tested (either in a validation set or with k fold cross validation); if performance does not meet a required threshold then more records are screened, chosen either through random sampling or, using active learning [10], on the basis either of those with highest uncertainty of predictions [11; 12] or alternatively from those most likely to be included[13; 14; 15]. Here we use a de novo search with subsequent training sets identified by random sampling, and we introduce a novel use of machine prediction, in identifying human error in screening decisions.

Machine learning approaches have been evaluated in context of systematic reviews of several medical problems including drug class efficacy assessment [7; 8; 12], genetic associations [9], public health [16; 13], cost-effectiveness analyses [9], toxicology [3], treatment effectiveness [17; 18] and nutrition [17]. To the best of our knowledge there have been only two attempts to apply such techniques to reviews of preclinical animal studies [3; 19]. These can be broad and shallow reviews or focussed and detailed reviews, and can have varying prevalence of inclusion.

Here we outline the ML approach taken to assist in screening a corpus for a broad and shallow systematic review seeking to summarise studies using non-human animal models of depression, based on a corpus of 70,365 records retrieved from two online biomedical databases. *In this paper, our aim was to identify the amount of training data required for an algorithm to achieve the level of performance of two independent human screeners, so that we might reduce the human resource required.*

Sena and colleagues developed guidelines for the appraisal of systematic reviews of animal studies [20]. These guidelines consider dual extraction by two independent human reviewers as a feature of a high quality review. From a large corpus of reviews conducted by CAMARADES we estimate the inter-screener agreement to be between 95% and 99%. Errors may occur at random (due to fatigue or distraction) or, more consequentially, systematic error, which, if included in a training set, might be propagated into a ML algorithm. Sources of systematic errors with certain types of records being at greater risk of misclassification. To our knowledge the nature of this 5% residual human error in systematic review methodology has not been formally investigated. The training data used for ML categorisation is based on training instances that has been screened by two independent human screeners.

*We therefore aimed to explore the use of established ML algorithms as part of a preclinical systematic review framework at the classification stage, to investigate if the ML algorithms could be used to improve the human gold standard by identifying human screening errors and thus improve the overall performance of ML.*

## Methods

We applied two independent machine learning approaches to the screening of a large (70,365 records) systematic review. Because we did could not predict how many training instances would be required we first selected 2000 records at random to provide the first training set. Of these, only 1993 were suitable due to data deposition errors. These were then screened by 2 human reviewers with previous experience with reviews of animal studies, with a third expert reviewer reconciling any differences. The resulting ML algorithms gave a score between 0 and 1. To ensure that the true sensitivity was likely to be 95% or higher we chose as our cut-point the value for which the lower bound of the 95% confidence interval of the observed sensitivity exceeded 95% when applied to the unseen validation dataset. We the repeated this process adding a further 1000 randomly selected (996 useable) citations to the training set; and then again adding a further 3000 randomly selected (2760 useable) citations to the training set. At each stage, performance of the approaches was assessed on a validation set of unseen documents, using a number of different metrics. Next, the best performing algorithm was used to identify human errors in the training and validation sets by selecting those with the largest discrepancy between the human decision (characterised as 0 for exclude or 1 for include) and the machine prediction (a continuous variable between 0 and 1). Performance of the approaches trained on the full 5749 records is reported here, and of each of the iterations is available in Supplementary Materials 1. The error analysis was assessed on the net reclassification index, and the performance of the ML approach is compared before and after correcting the errors in human screening using AUC.

### Step 1: Application of ML tools to screening of a large preclinical systematic review

#### Training Sets

70,365 potentially relevant records were identified from Pubmed and EMBASE The search strings were composed of the animal filters devised by the Systematic Review Center for Laboratory animal Experimentation (SYRCLE) [21; 22], NOT reviews, comments, or letters AND a depression disorder string (for full search strings see [23]). The training set and the validation set were chosen at random from the 70,365 by assigning each record a random number using the RAND function in excel and ranking them from smallest to largest. The training set consisted of 5749 records. The validation set consisted of 1251 records. The training set and validation set were screened by two independent human screeners with any discrepancies reconciled by a third independent human screener. The human screening process involved an online tool (app.syrf.org), which randomly presents a reviewer with a record, with the title and abstract displayed. The reviewer makes a decision about the record, included (1) or excluded (0). A second reviewer is also randomly presented with records. If a record receives two ‘included’ decisions, the screening for this record is considered complete. If reviewer 1 and reviewer 2 disagree, the record gets presented to a third reviewer who makes a decision. The record then has an average inclusion score of 0.666 or 0.333. Any record that has an inclusion score above 0.6 is included, those scoring less than 0.6 are excluded, and screening is considered complete. Datasets are available on Zenodo, as described in “Availability of Data & Materials” below, Performance was assessed at each level on a validation set of unseen records. The training and validation set were selected consecutively from the initial random ordering. For the training set of 5749 records, the validation set was the subsequent 1251 records. This validation set had more than 150 “included” records, which can give reasonably precise 95% confidence intervals for sensitivity and specificity.

**Figure 1.**
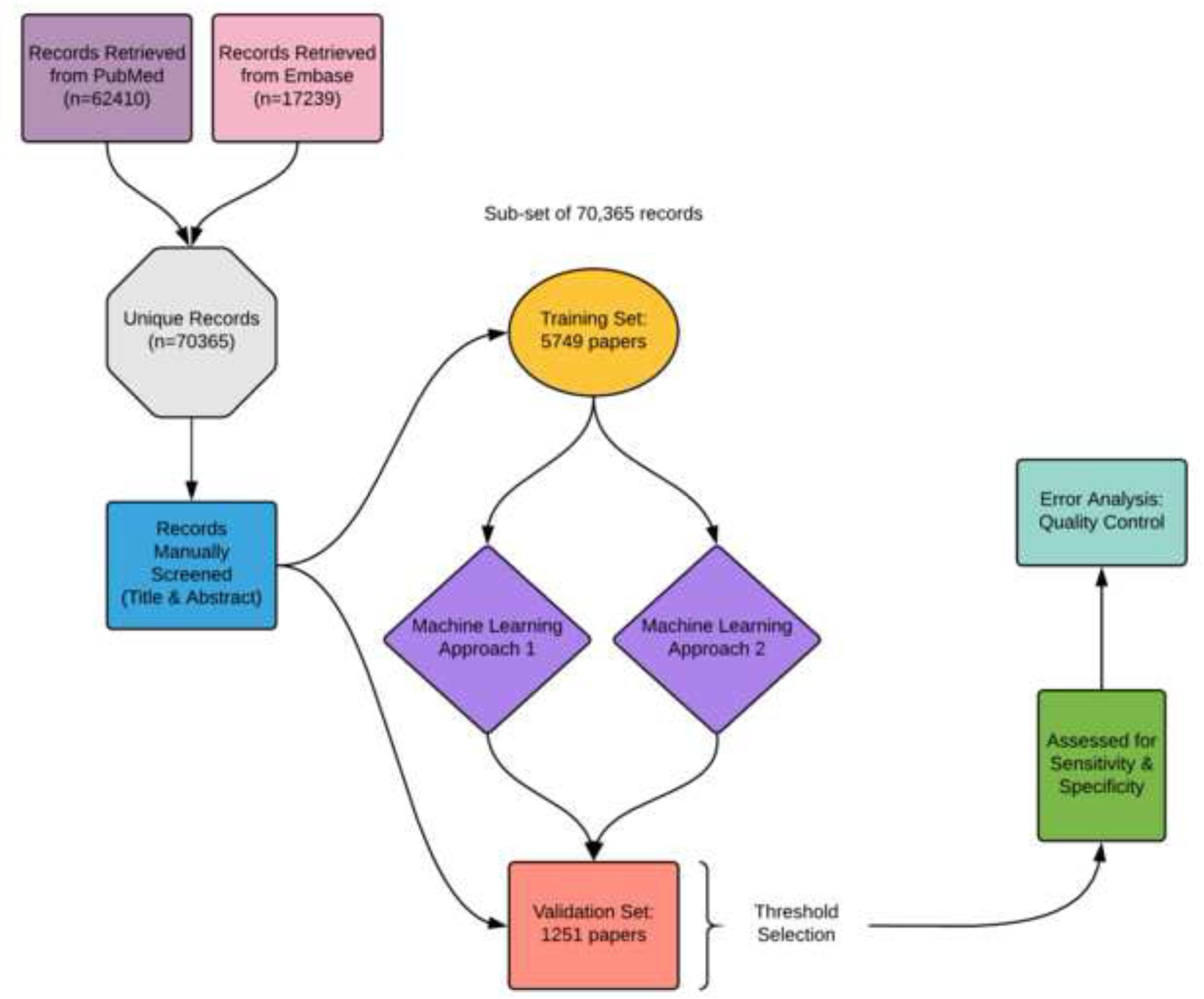
Diagram of the Layout of the Study.

#### Feature Generation

First, documents in the training set were transformed into a representation appropriate for the machine learning algorithms. Documents were created by concatenating the title and the abstract. Every case (document) is represented by a fixed number of features, numerical quantities describing certain properties that might be used by the classifier to extract rules and make predictions about inclusion. The classifiers described below used generally similar approaches

We used “bag-of-words” (BoW) to characterise document titles and abstracts in both classifiers. To account for the relative importance of words within a given document, and difference in words used between documents we used ‘Term Frequency – Inverse Document Frequency’ (TD-IDF). This is defined as:

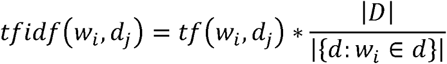

The score for the i-th word in context of the j-th document takes into account not only how many times the word occurred there (tf), but also how many other documents (d) from the whole corpus (D) contain it as well. This helps to reduce the score for words that are common for all documents and therefore have little predictive power. This helps the classifier to focus on terms which help to distinguish between documents, rather than on terms which occur frequently [24]. We allowed n-grams; did not use stemming; and used the MySQL text indexing functionality “stopword” list to remove frequently occurring words which provide little relevant information for classification purposes. [25]

Because bag-of-words representation generates as many unigram features as there are words in the collection (typically at least several thousand); and many more when using higher-order n-grams, we used additional approaches. Latent Semantic Indexing (LSI) and Latent Dirichlet Allocation (LDA) represent textual data in a more efficient way. In LSI [26], the training set is represented as a matrix, where rows correspond to documents, columns to terms (words or n-grams), while cells contain frequency or TF/IDF score of a given term in a given document. The matrix is then decomposed using a general matrix factorisation technique known as Singular Value Decomposition (SVD) and truncated to the first n dimensions. Because of the properties of SVD the new features will be such linear combinations of features of the old space that minimise the differences between the original and the transformed space. In case of textual data it means that those words that frequently occur in the same documents (probably because of the similar meaning) will be treated in the same way. The n is set a-priori to a reasonably low value – usually a few hundred. LDA exploits distributional similarities between words, but based on explaining document contents using a Bayesian network [27]. This method is based on the premise that every document is a mixture of topics, which in turn consist of related words. The correspondence between documents and topics and between topics and words can be inferred via Gibbs sampling process. As a result, similarly to LSI, every document is represented by a sequence of n numbers, indicating how related it is to every topic [28]. Unlike in SVD, the model fitness to the data cannot be expressed through the amount of variance of the original matrix it explains and the optimal number of topics may be different for every collection and classification task. Following previous work in the domain [13] and the user guide for MALLET (the tool we use for LDA, which recommends values between 200 and 400) we elected to generate 300 topics Here we use three feature sets, BoW, LDA and SVD (LSI) individually, in pairs and finally all together; preliminary evaluation through the cross-validation on the training set suggests that LDA+SVD and bag-of-words with a simple linear classifier deliver the most robust performance.

#### Classifiers

Following the transformations made in feature selection, the documents are then used to train the machine learning classifier. The classifier most commonly used for document classification in context of systematic reviews [11; 13; 8; 9; 12; 14; 15; 17] is the Support Vector Machine (SVM) as it has frequently been used for tasks involving text.). SVM is a supervised learning algorithm, learning to classify new documents based on a training set of labelled documents [31]. This algorithm represents training documents as points in a multi-dimensional space defined by all available features. To be able to classify cases into positive and negative category, it seeks a hyperplane dividing the space into one side corresponding to included documents and the other to excluded ones. Based on the training data, the optimal hyperplane is constructed so that it maximises both the number of training cases located on the “correct” side of decision boundary and their distance from the plane (margin). The new, unseen, documents are then ranked according to their location with respect to the boundary. Those far from it are confidently predicted as included or excluded, according to which side of the plane they lie. The cases which the model has less confidence about will be located close to the hyperplane. Logistic regression is a similar linear classifier, which instead of hyperplane, seeks such coefficients of a linear combination of feature values that will give high values for positive cases (included documents) and low for negative (excluded documents). Both of these approaches could be enriched with feature selection elements to mitigate the problems with multitude of features.

Three feature sets (BoW, LDA and SVD (LSI)) were tested on SVMs, logistic regression and random forests [32]. The two algorithms described below performed best for this dataset of 70,365 records, on the broad topic of preclinical animal models of depression.

#### Approaches

Here, two approaches were developed independently, using different classification models and feature representations, but sharing the linear classification principles.

##### Approach 1

Approach one used a tri-gram ‘bag-of-words’ model for feature selection and implemented a linear support vector machine with Stochastic Gradient Descent (SGD) as supported by the SciKit-Learn python library [33]. This classifier was chosen it is efficient, scales well to large numbers of records, and provides an easily interpretable list of probability estimates when predicting class membership (i.e. scores for each document lying between 0 and 1). Efficiency and interpretability are important, as this classifier is already deployed in a large systematic review platform [34], and any deployed algorithm therefore needs not to be too computationally demanding, and its results understood by users who are not machine learning specialists. The tri-gram feature selection approach without any additional feature engineering also reflects the generalist need of deployment on a platform used in a wide range of reviews: the algorithm needs to be generalisable across disciplines and literatures, and not ‘over-fitted’ to a specific area. For example, the tri-gram “randomised controlled trial” has quite different implications for classification compared with “randomised controlled trials” (i.e. ‘trials’ in plural). The former might be a report of a randomised controlled trial; while the latter is often found in reports of systematic reviews of randomised trials. Stemming would remove the ‘s’ on trials and thus lose this important information. Here, the algorithm needs to be generalisable across disciplines and literatures, and not be ‘over-fitted’ to a specific area. This approach aims to give the best compromise between reliable performance across a wide range of domains and that achievable from a workflow that has been highly tuned to a specific context.

##### Approach 2

Approach 2 used a regularised logistic regression model built on LDA and SVD features. Namely, the document text (consisting of title and abstract) was first lemmatised with the tool GENIA tagger [35] and then converted into bag of words representation of unigrams, which was then used to create two types of features. First, the word frequencies were converted into a matrix TF/IDF scores, which was then decomposed via SVD implemented in scikit-learn library and truncated to the first 300 dimensions. Second, an LDA model was built using MALLET library [36], setting 300 as a number of topics. As a result each document was represented by 600 features, and an L1-regularised logistic regression model was built using glmnet package [37] in R statistical framework [38].

In this procedure every document is represented with a constant, manageable number of features, irrespective of corpus or vocabulary size. As a result, we can use a relatively simple classification algorithm and expect good performance with short processing time even for very large collections. This feature is particularly useful when running the procedure numerous times in cross-validation mode for error analysis (see below).

For a given unseen test instance, the logistic regression returns a score corresponding to the probability of it being relevant according to the current model. An optimal cut-off score that gives the best performance is calculated as described above.

### Assessing Machine Learning Performance

The facets of a machine learning algorithm performance that would be most beneficial to this field of research are high sensitivity (see table 1), at a level comparable to the 95% we estimate is achieved by two independent human screeners. We therefore need to be confident that the sensitivity is 95% or higher, which we do by setting our cut point such that the lower bound of the 95% confidence interval of the observed sensitivity is 95% or higher. Once the level of sensitivity has been reached, the aim is to maximise specificity, to reduce the number of irrelevant records included by an algorithm. Although specificity at 95% sensitivity is our goal, we provide values of other measures for better illustration of the performance.

#### Performance metrics

Performance was assessed using sensitivity (or recall), specificity, precision, accuracy, and Work Saved over Sampling (WSS) (see table 1), carried out in R (R version 3.4.2; [38]) using the ‘caret’ package [39]. 95% Confidence Intervals were calculated using the efficient-score method [40]. Cut-offs for were determined manually for each approach by taking the score that achieved 95% sensitivity (including the lower 95% confidence level), and the specificity at this score was calculated.

**Table 1.**
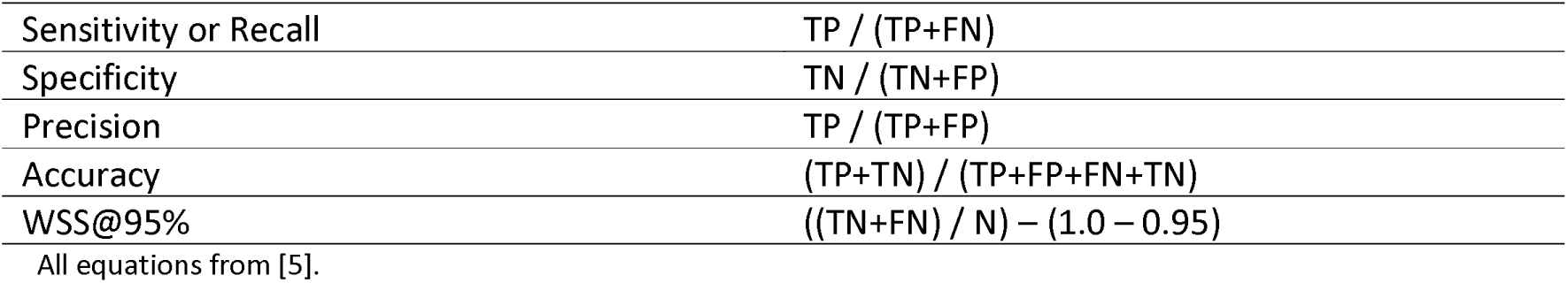
Equations used to assess performance of machine learning algorithms

### Step 2: Application of ML tools to training datasets to identify human error

#### Error Analysis Methods

The methodology for the error analysis was outlined in an *a priori* protocol, published on the CAMARADES website on 18^th^ December 2016 [41]. To generate the machine learning scores for the set of records that were originally used to train the machine (5749 records), the non-exhaustive cross-validation method, 5-fold validation, was used. This method involved randomly partitioning the set of records into 5 equal sized subsamples. One subsample was set aside, and the remaining 4 subsamples were used to train the algorithm [42]. Thanks to this process, every record has a score computed by a machine learning model built without including it in the training portion. These scores were used to highlight discrepancies or disagreements between machine decision and human decision. The documents were ordered by the machine assigned labels in order of predictive probability, from most likely to be relevant to least likely to be relevant. The original human assigned scores were placed next to the machine-assigned scores, to highlight potential errors in the human decision. A single human reviewer (experienced in animal systematic reviews) manually reassessed the records where discrepancies were highlighted starting with the most discrepant. To avoid reassessing the full 5749 record dataset, a stopping rule was established such that if the initial human decision was correct for five consecutive records, further records were not reassessed.

**Figure 2.**
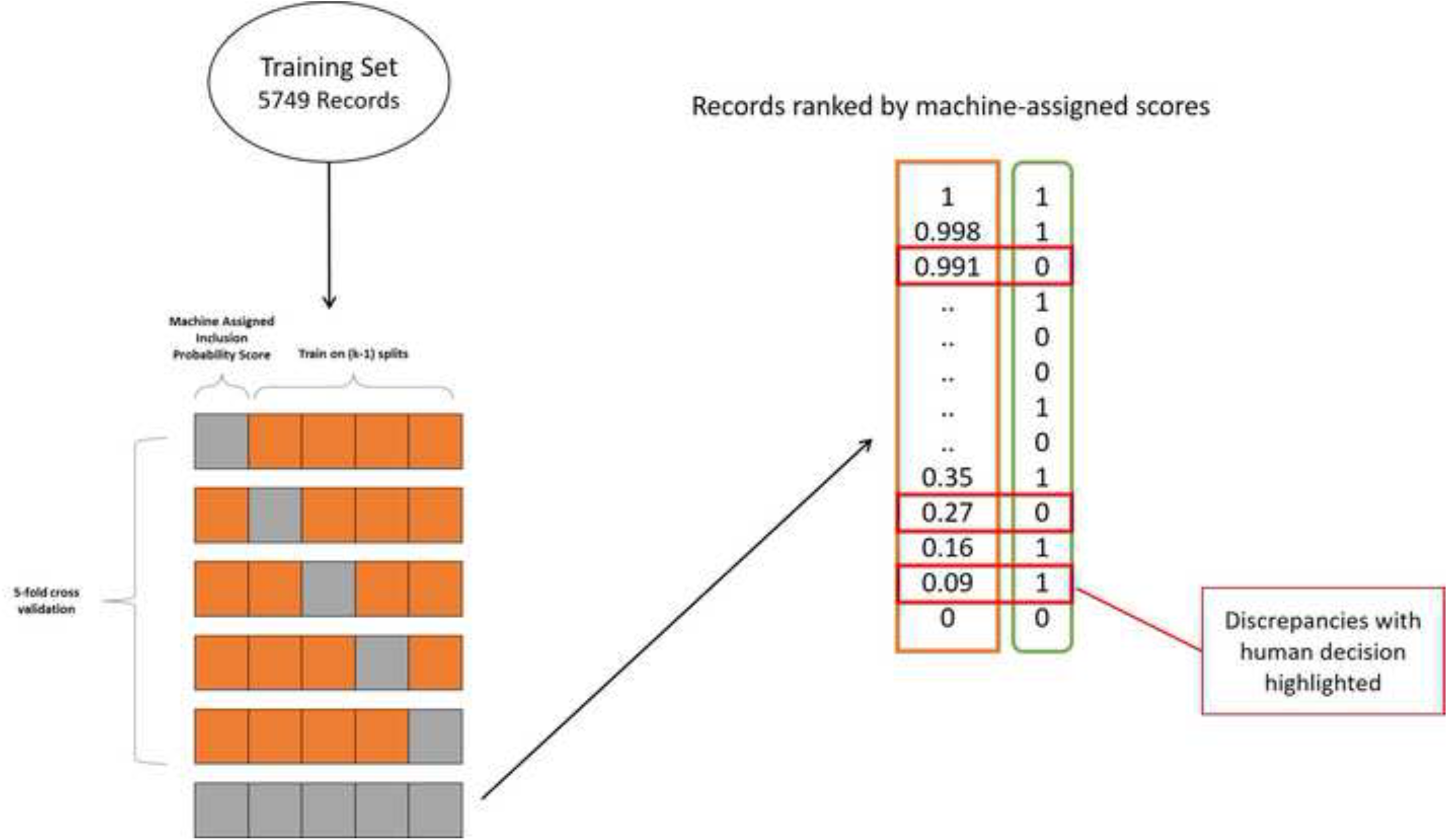
Error Analysis. *The methodology for using cross-validation to assign ML predicted probability scores. The ML predicted probability scores for the records were checked against the original human inclusion decision.*

After the errors in the training set were investigated and corrected as described above a new model was built on the updated training data. The outcome of error analysis is presented as reclassification tables, the area under the curve (AUC) being used to compare the performance of the ML algorithm trained on the ‘old’ training set of records, and the net reclassification index (NRI) [43] used to compare the performance of the classifier built on the updated training data with the performance of the classifier built on the original training data. The following equation was used:

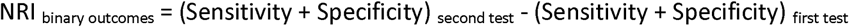
[44]

The AUC was calculated using the DeLong method in the ‘pROC’ package in R [45].

Further, we applied the same technique as above to identify human screening errors in the validation dataset. Due to the small number of records in the validation set (1251 records), it was assumed that every error would be likely to impact measured performance, and so the manual screening of the validation set involved revisiting every record where the human and machine decision were incongruent. The number of reclassified records was noted. The inter-rater reliability of all screening decisions on training set and validation set between Reviewer 1 and Reviewer 2 were analysed using the ‘Kappa.test’ function in the ‘fmsb’ package in R [46].

## Results

In this section we first describe the performance from the ML algorithms. We then show the results from the analysis of human error, and finally describe the performance of the ML algorithm after human errors in the training and validation set have been corrected.

### Performance of Machine Learning Algorithms

Table 2 shows the performance of the two machine learning approaches from the SLIM (Systematic Living Information Machine) collaboration. The desired sensitivity of 95% (including lower bound 95% CI) has been reach by both approaches. Both approaches reached 98.7% sensitivity based on learning from a training set of 5749 records, with an inclusion prevalence of 13.2% (see below). Approach 1 reached a higher specificity level of 86%. This is visualised on an AUC curve (figure 1).

**Table 2.**
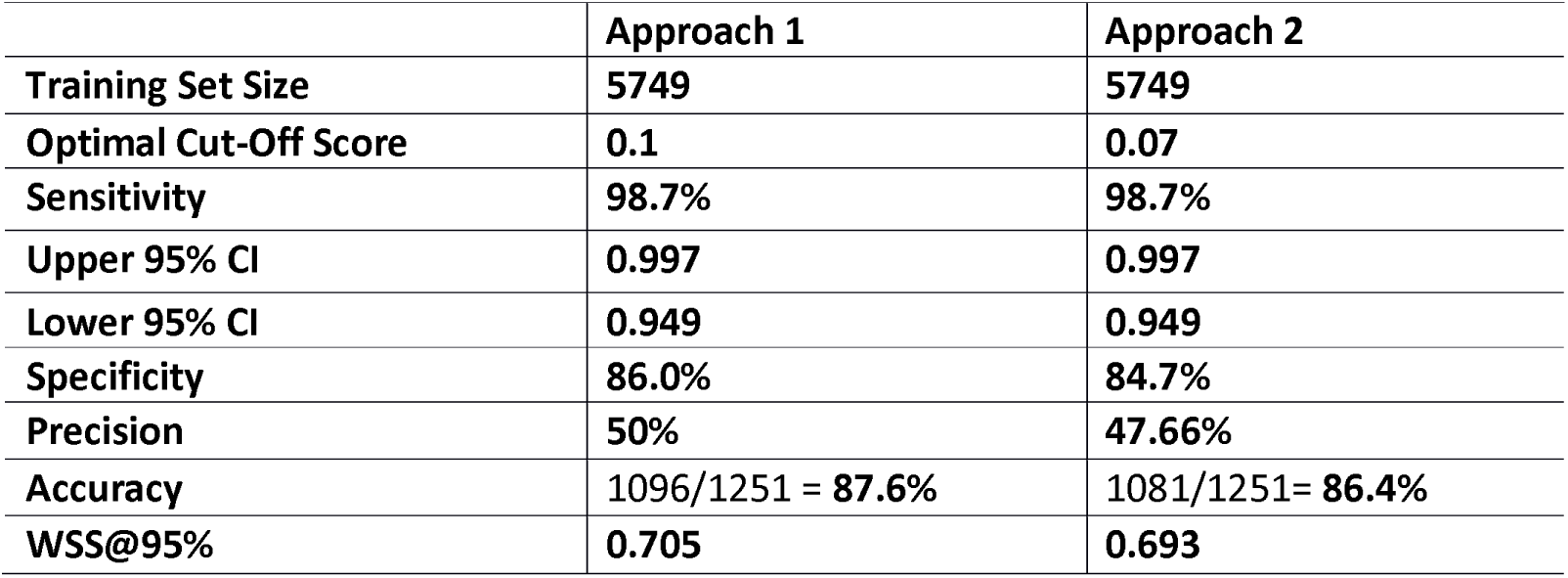
Performance of machine learning approaches on depression training dataset.

**Figure 3.**
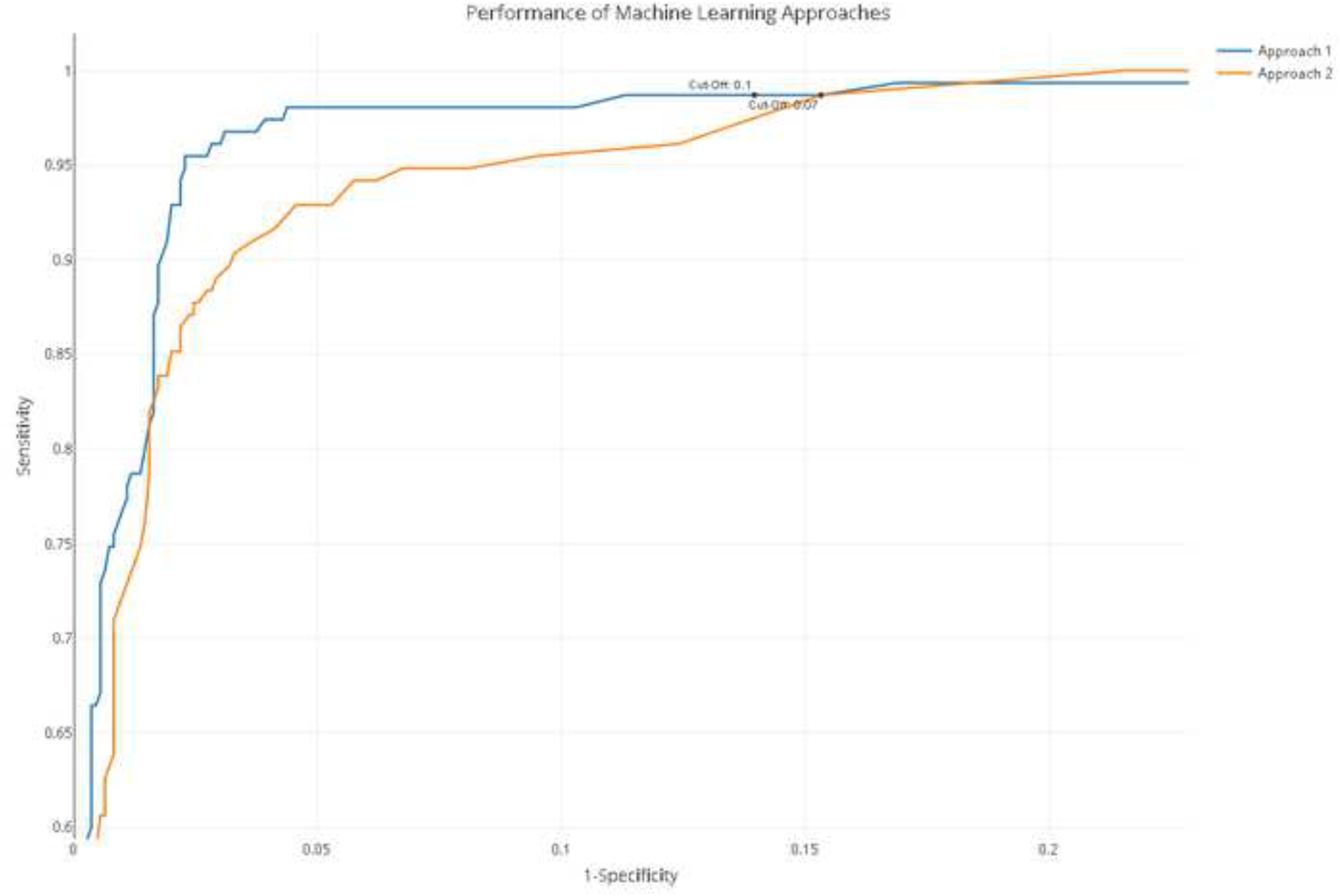
Performance of Machine Learning Approaches. *For the interactive version of this plot with cut-off values, see code and data at https://github.com/abannachbrown/The-use-of-text-mining-and-machine-learning-algorithms-in-systematic-reviews/blob/master/ML-fig3.html*

### Error Analysis & Reclassification

Cohen’s κ was run to determine the interrater agreement of screening decisions between Reviewer 1 and Reviewer 2. Κ = 0.791 (95% CI, 0.769 to 0.811), p < 0.0001, with 281 records requiring a third reviewer decision. To assess whether machine learning algorithms can identify human error and therefore improve the training data, error analysis was conducted. Seventy-five papers out of 5749 papers had predictive scores very far from the human assigned labels, so were reassessed to see if these were due to human errors. Out of 75 rescreened papers, the machine corrected the human decision 47 times. The machine was wrong, (i.e. the initial human decision was correct) 28 times. The validation set was also rescreened. Ten papers out of the 1251 records were identified as potential human errors. Out of 10 errors, the machine corrected 8 human decisions. These 8 records were all falsely excluded by the human and were now included. The initial human decision was correct twice.

To calculate human error in the training set, the number of errors identified (47) out of the training set (5749 records) was calculated to be at least 0.8%. Of the 47 records reclassified, 11 records were falsely included in the original screening process and were now correctly excluded, and 36 records were falsely excluded in the original screening process and were now correctly included. The machine correctly identified human screening errors, which were calculated to be just under 1% of the dual screened training set. Forty-seven papers out of 760 were ‘correctly’ reclassified, 6% of the included papers.

Similarly, the human error rate in the validation set (1251 records) was 0.6%. Again looking at the prevalence of inclusion in this dataset (155/1251), which is 12.4%, the 8 records of out the now 163 were correctly reclassified which is 4.9% reclassified. All 8 records we falsely excluded in the original screening process and are now correctly included.

Test 1: 98.7% + 86% = 184.7%

Test 2: 98.2% + 89.3% = 187.5%

**NRI = 3.2%**

We consider the updated validation set to be the new gold standard as 8 records were now included. The confusion matrix for the performance of the machine learning algorithm after the error analysis update on the training records is displayed below in table 3.

**Table 3.**
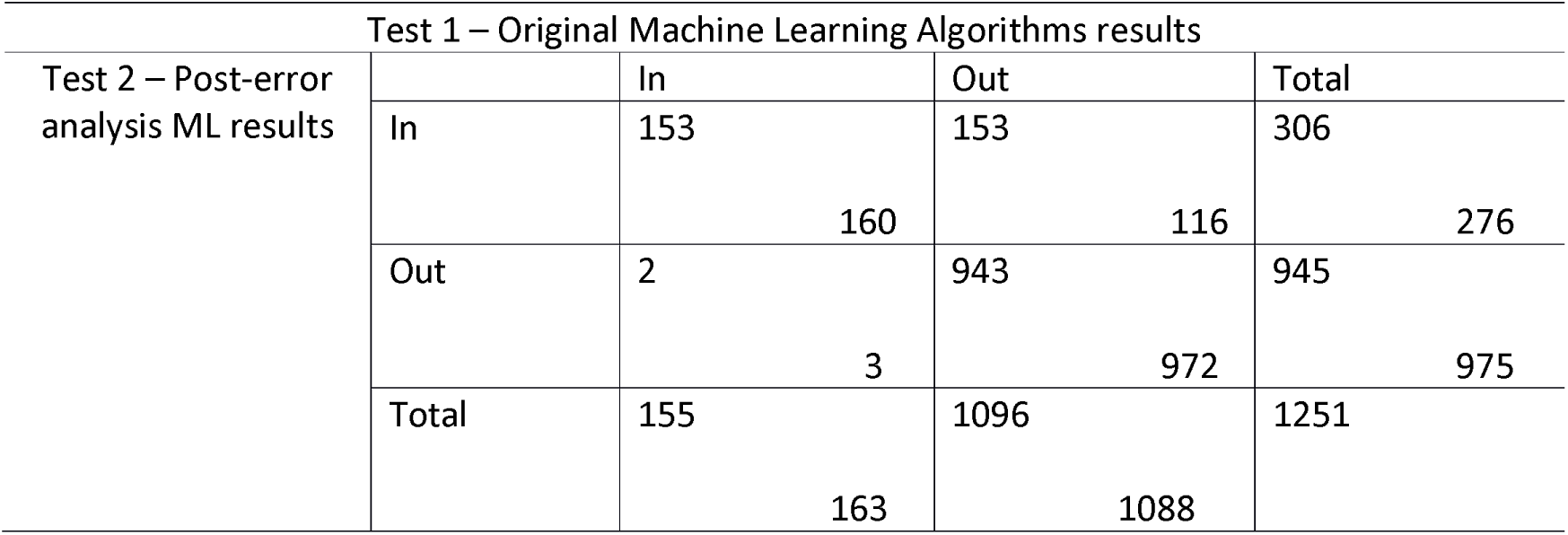
Reclassification of records in validation after error analysis.

Analysing the human errors identified by the machine learning algorithm and correcting for these errors and re-teaching the algorithm leads to improved performance of the algorithm, particularly its sensitivity. This can save considerable human time in the screening stage of a systematic review. Consider the remaining approximately 64,000 papers, if the ML algorithm results are 3% more accurate, that is approximately 2000 papers that are correctly ‘excluded’ that would not be forwarded for data extraction.

### After Error Analysis: Improving Machine Learning

Using the error analysis technique above, of the 47 errors identified in the full training dataset of 5749 records, 0.8% were corrected. We retrained approach 1 on the corrected training set and measured performance on the corrected validation set of 1251 records as we consider this to be the ‘new’ gold standard. The performance of the original approach 1 and updated approach 1 was assessed on the corrected validation set of 1251 records. The performance of this retrained algorithm in comparison to the performance of the original classifier 5 on the updated validation set is shown in table 4.

**Table 4.**
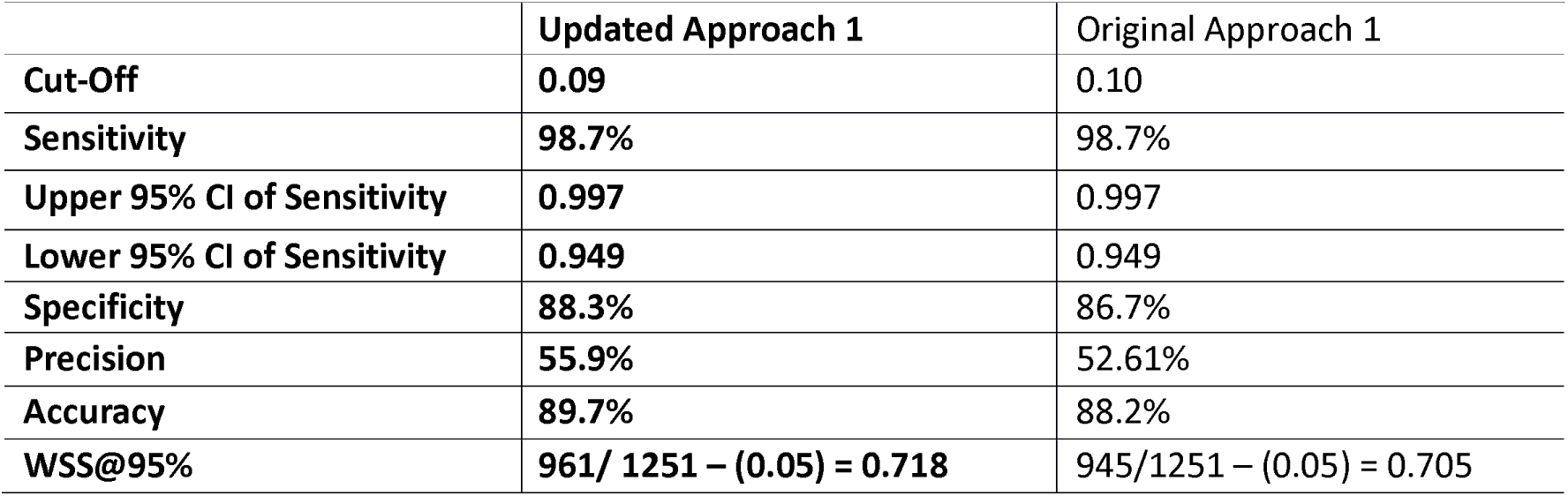
Performance of machine learning approach after error analysis.

**Figure 4.**
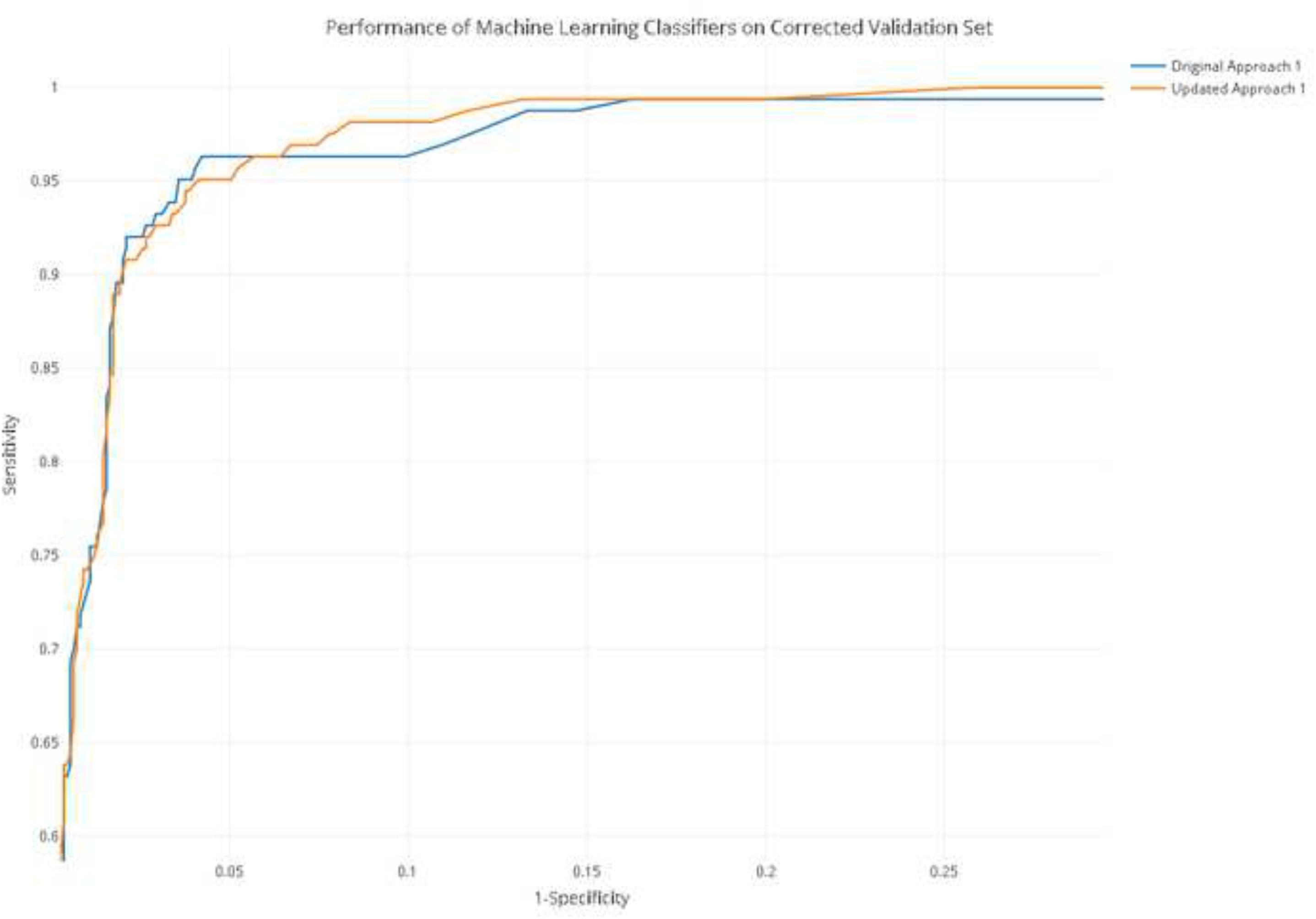
Performance of Approach 1 after error analysis. *The updated approach is retrained on the corrected training set after error analysis correction. Performance on both the original and the updated approach is measured on the corrected validation set (with error analysis correction). For the interactive version of this plot with exact cut-off values, see code and data at https://github.com/abannachbrown/The-use-of-text-mining-and-machine-learning-algorithms-in-systematic-reviews/blob/master/error-analysis-plot.html*

We compared the area under the ROC curve for the original approach 1 and the updated approach 1. The AUC for the original approach 1 was 0.9272 (95% CI calculated using DeLong method; 0.914-0.9404). The AUC for the updated approach 1 was 0.9355 (95% CI calculated using DeLong method; 0.9227-0.9483). DeLong’s test to compare the AUC between the ROC of the two approaches was applied ‘, Z = −2.3685, p = 0.0178.

## Discussion

### Document Classification

We have shown machine learning algorithms to have high levels of performance, with 98.7% sensitivity and 88.3% specificity; this sensitivity is comparable to two independent human screeners. The objectives for selecting ML approaches in this project was to achieve a minimum 95% sensitivity (including lower bound confidence intervals), to minimise the number of potentially relevant papers which are wrongly excluded. Thereafter, algorithms were then chosen on the basis of their specificity. to reduce the subsequent human time required to sort through and assess papers.

The two approaches have similar performance. The slight differences may reflect the method of feature generation. These algorithms have high performance on this specific topic of animal models of depression. As demonstrated previously, the performance of various classifiers can alter depending on the topic and specificity of the research question [3].

In this study, the cut-off points were selected using the decisions on the validation set to achieve the desired performance. Although this allows the measurement of the maximum possible gain using a given approach in an evaluation setting, in practice (e.g. when updating a review), the true scores would not be available. The problem of choosing a cut-off threshold, equivalent to deciding when to stop when using a model for prioritising relevant documents, remains an open research question in information retrieval. Based on their experience with a given tool, a reviewer may come up with a heuristic fitting their workflow, e.g. if no new includes are seen in the 100 highest-ranked documents, then everything else could be discarded as well. More sophisticated approaches have also been tested [47], but they do not guarantee achieving a desired sensitivity level. It has to be noted that ML-based prioritisation could be useful even if no cut-off is used and all documents are screened manually, since seeing the relevant documents first can help to organise the process and thus reduce the workload [5]. In a similar broad preclinical research project in neuropathic pain it took 18 person months to screen 33,814 unique records – based on these numbers it would take an estimated 40 person months to screen 70,365 unique records. Performance of machine learning tools demonstrated in this paper can greatly reduce the amount of human resource needed for initial title and abstract screening of a large corpus of records retrieved from a broad search.

We have applied the algorithm to the full dataset (remaining 63,365 records) and are in the process of full-text screening. Following this process, it will allow a more in depth learning on the part of the machine that it can apply to any updates to the search.

### Error Analysis

By using the ML algorithm to classify the likelihood of inclusion for each record in the training set, we highlighted discrepancies between the human inclusion or exclusion decision and the machine decision. Using this technique, we identified human errors, which were then corrected to update the training set.

Human screening of the training set was conducted using the “majority vote” system; it is interesting to consider the potential reasons for errors or ‘misclassifications’ arising in this process. Reviewers’ interpretation of the “breadth” of this wide review might be one contributing factor to discrepancies. With a less clear cut-off, reviewers are unsure of where some articles should be included. Discrepancies arising where Reviewer 1 was more inclusive and where Reviewer 2 was less inclusive, thereby Reviewer 3 will be the deciding factor. A different approach whereby Reviewer 1 and 2 discuss discrepancies might be a pinpoint the exact reasons for misunderstandings or different interpretations of the inclusion criteria. However, for larger projects when using a crowd-sourcing approach with many individual people contributing to each Reviewer, this may not be a practical solution.

We have successfully identified human screening errors which were calculated to be just under 1% of the training set which was dual screened by two independent human reviewers. The prevalence of inclusion in this training set is 13.2% (760 out of the 5749), so an error of 0.8% is likely to be important Therefore errors of false inclusion or exclusion in the training sets may have a substantial impact on the learning of the ML algorithm. This error analysis results in a 3% increase or change in sensitivity and specificity, with increased precision, accuracy, and work saved over sampling of the algorithm. We observed an increase in specificity of 1.6% without compromise to sensitivity. In a systematic review with this number of records this saves considerable human resources, as the number of records required to screen reduces by at least 1125.

This error analysis was an initial pilot with stopping criteria where if the initial human decision was correct five consecutive times, further records were not reassessed. It is possible and likely that there are further errors in the human screened training set. A more in-depth analysis of the training dataset, investigating every instance where the human and machine decision were incongruent, might identify more errors and further increase the precision and accuracy of machine learning approaches, further reducing human resources required for this stage of systematic review. We have shown here that even with minimal intervention (only assessing incongruent records until the original human decision was correct 5 consecutive times), the performance of ML approaches can be improved.

### Limitations & Future Directions

Here we show the best performing algorithms for this dataset with a broad research question. Other dissimilar research questions or topics may require different levels of training data to achieve the same levels of performance, or may require different topic modelling approaches or classifiers. The best performing algorithm, outlined in this paper, is being applied in an ongoing research project, therefore the ‘true’ inclusion and exclusion results for the remaining 63365 records is not yet known. The ‘true’ results will unfold with the fullness of time.

These machine learning algorithms are deployed in an existing systematic review online platform, EPPI-Reviewer [34], and are in the process of being integrated into the Systematic Review Facility (SyRF) tool, which is focused on the preclinical domain (www.app.syrf.org). This will improve the ease of use of machine learning functions for systematic reviewers, increase the usage of machine learning algorithms for systematic review and significantly reduce the amount of human resources required to conduct systematic review across a range of topics. By allowing a degree of user control over which classifiers and the levels of performance are required for each specific research project. With a broad collaboration such as SLIM we aim to test many ML algorithms across a range of research topics to identify which classifiers perform best under which circumstances, to be able to provide recommendations to users of SyRF.

This paper outlines a pilot approach to using machine learning algorithms to identify human errors in current systematic review methodology. Future research can investigate this concept more thoroughly by setting up a more comprehensive experimental design. After further investigation into the extent of human error in dual reviewing, the picture will be clearer as to the scale of human error and to what extent a machine learning algorithm can identify and aid in rectifying this. These tools can could be integrated into systematic review platforms, such as SyRF (www.app.syrf.org), and may provide feedback to the systematic reviewer during screening, and could ultimately flag incorrectly screened records as the human screens them for inclusion in a dataset for machine training.

### Conclusions

We have demonstrated that machine learning techniques can be successfully applied to an ongoing, broad pre-clinical systematic review. We have demonstrated that machine learning techniques can be used to identify human errors in the training and validation datasets. We have demonstrated that updating the learning of the algorithm after error analysis improves performance. This error analysis technique requires further detailed elucidation and validation. These machine learning techniques are in the process of being integrated into existing systematic review applications to enable more wide-spread use. In future, machine learning and error analysis techniques that are optimised for different types of review topics and research questions can be applied seamlessly within the existing methodological framework.

## List of Abbreviations

AUC: Area Under the Curve
BoW: Bag-of-Words
CAMARADES: Collaborative Approach to Meta-Analysis and Review of Animal Data from Experimental Studies
LDA: Latent Dirichlet Allocation
LSI: Latent Semantic Indexing
ML: Machine learning
NRI: Net Reclassification Index
PROSPERO: International Prospective Register of Systematic Reviews
SVD: Singular Value Decomposition
SLIM: (Systematic Living Information Machine) collaboration
SGD: Stochastic Gradient Descent
SVM: Support Vector Machine
SYRCLE: Systematic Review Center for Laboratory animal Experimentation
SyRF: Systematic Review Facility
TD-IDF: Term Frequency – Inverse Document Frequency
WSS: Work Saved over Sampling

## Declarations

### Ethical Approval

Not applicable.

### Availability of Data & Materials

The training and validation datasets, error analysis datasheets, as well as all the records in the depression systematic review are available on Zenodo: DOI 10.5281/zenodo.60269

The protocol for the systematic review of animal models of depression is available from: http://onlinelibrary.wiley.com/doi/10.1002/ebm2.24/pdf

The protocol for the Error Analysis is available via the CAMARADES website and can be accessed directly from this link: https://drive.google.com/file/d/0BxckMffc78BYTm0tUzJJZkc1alk/view

The results of the classification algorithms and the R code used to generate the results is available on GitHub: https://github.com/abannachbrown/The-use-of-text-mining-and-machine-learning-algorithms-in-systematic-reviews.

### Authors’ Contributions

ABB screened and analysed the datasets. JT & PB conducted feature selection and built the classifiers. ABB, JT & PB wrote the manuscript. ABB, JT, PB, MRM, JL, AR & SA devised the study. JL, MRM & SA supervised the study. All authors edited and approved the final manuscript.

## Acknowledgements

Thank you to Kaitlyn Hair, Paula Grill, Monica Dingwall & Zsanett Bahor for their assistance in second screening the training and validation datasets.

